# NOTCH inhibition promotes myoblast fusion by releasing HEYL repression on *TMEM8C* regulatory regions in foetal skeletal muscles

**DOI:** 10.1101/2020.01.09.900159

**Authors:** Joana Esteves de Lima, Cédrine Blavet, Marie-Ange Bonnin, Estelle Hirsinger, Emmanuelle Havis, Delphine Duprez

**Affiliations:** Sorbonne Université, Institut Biologie Paris Seine, CNRS UMR7622, Developmental Biology Laboratory, Inserm U1156, F-75005 Paris, France; Biology of the Neuromuscular System, INSERM IMRB U955-E10, UPEC, ENVA, EFS, Creteil 94000, France

**Keywords:** NOTCH, HEYL, TMEM8C, MYOG, myoblast fusion, foetal myogenesis, limb, chicken

## Abstract

Differentiation and fusion are two intricate processes involved in skeletal muscle development. The close association of differentiation and fusion makes it difficult to address the process of fusion independently of differentiation. Using the fusion marker *myomaker*, named *TMEM8C* in chicken, we found that both *TMEM8C* transcripts and the differentiated and fusion-competent MYOG+ cells are preferentially regionalized in the central regions of limb foetal muscles in chicken embryos. Because the NOTCH signalling pathway is a potent inhibitor of muscle differentiation during developmental myogenesis, NOTCH function in myoblast fusion was not addressed so far. We analysed the consequences of NOTCH inhibition for myoblast fusion and *TMEM8C* expression during foetal myogenesis using in vitro and in vivo chicken systems. NOTCH inhibition following chicken embryo immobilisation or in myoblast cultures increased *TMEM8C* expression and myoblast fusion. Moreover, we showed that NOTCH inhibition induced the un-binding of the HEYL transcriptional repressor from the *TMEM8C* regulatory regions in limb muscles and myoblast cultures. These results identify a molecular mechanism underlying the fusion-promoting effect of NOTCH-inhibition during foetal myogenesis.

## Introduction

Skeletal muscle development, homeostasis and regeneration rely on muscle stem cells that undergo a multistep process to form multinucleated cells named myofibres. The myofibres, the main structural unit of skeletal muscle, are formed by myoblast fusion. The fusion process has been poorly studied because it is difficult to dissociate the differentiation and fusion processes during myogenesis.

The transcriptional control of the skeletal muscle programme is dependent on four bHLH transcription factors, MYF5, MYOD, MRF4 and MYOG, named the Myogenic Regulatory Factors (MRFs) (Buckingham and Rigby, 2014). MRFs have the ability to trigger the muscle program including the successive steps of specification, differentiation and fusion from muscle progenitors (Buckingham and Rigby, 2014; Comai and Tajbakhsh, 2014), but also from non-muscle cells in vitro (Weintraub et al., 1991) and in vivo (Delfini and Duprez, 2004). Multinucleated myofibre formation is a multistep process involving cell cycle withdrawal of already specified myoblasts, cell elongation, cell-cell contact and fusion (Biressi et al., 2007; Comai and Tajbakhsh, 2014). The three main steps that underlie myoblast fusion are: (1) cell recognition and adhesion; (2) enhancement of cell proximity via F-actin-propelled membrane protrusions from one fusion partner cell and myosin II-dependent cortical tension in the other fusion partner cell; (3) destabilization of the two apposed plasma membrane lipid bilayers and formation of a fusion pore (Hernández and Podbilewicz, 2017; Kim et al., 2015). Numerous transmembrane proteins and molecules of the intracellular actin machinery are involved in myoblast fusion in vertebrates. However, they are not specific to myoblast fusion and are recruited for any cell-cell fusion processes such as sperm-egg fusion in fertilisation, cytotrophoblast fusion in placentation and axonal fusion during neuronal repair (Hernández and Podbilewicz, 2017). Recently, a muscle-specific gene, *myomaker* (named *TMEM8C* in chicken, *Mymk* in mice and *MYMK* in humans) has been identified as being essential for myoblast fusion in mice, chicken and zebrafish during development and muscle regeneration (Landemaine et al., 2014; Luo et al., 2015; Millay et al., 2013, 2014, 2016). Myomaker is a transmembrane protein of 221 amino acids with seven membrane-spanning regions. Autosomal recessive mutations in the *MYMK* gene, which reduce but do not eliminate *MYMK* function, cause a congenital myopathy, the Carey-Fineman-Ziter syndrome, in humans (Di Gioia et al., 2017). Myomaker is sufficient to trigger fibroblast-myoblast fusion but not fibroblast-fibroblast fusion (Millay et al., 2014). However, when combined with the micropeptide myomixer (also named myomerger or minion), myomaker triggers fibroblast-fibroblast fusion (Bi et al., 2017; Quinn et al., 2017; Zhang et al., 2017). By itself, myomixer does not possess 10T1/2 fibroblast-C2C12 myoblast fusion activity (Bi et al., 2017). Two MRFs, MYOD and MYOG, have been shown to positively regulate the transcription of *myomaker* genes in mouse, zebrafish and chicken via E-boxes located in the *myomaker* regulatory regions (Ganassi et al., 2018; Luo et al., 2015; Millay et al., 2014). The direct transcriptional regulation of the *myomaker* gene by MRFs complicates the distinction between the differentiation and fusion processes.

The identification of myomaker as a transmembrane protein required for myoblast fusion (Millay et al., 2013) stimulated the research on myoblast fusion. However, the signalling pathways that regulate *myomaker* expression and muscle fusion are not fully identified. NOTCH signalling is recognized to being involved in skeletal muscle differentiation during development. NOTCH inhibition promotes muscle differentiation in both in vivo and in vitro systems (Kitzmann et al., 2006; Kopan et al., 1994; Schuster-Gossler et al., 2007; Vasyutina et al., 2007). Conversely, NOTCH is a potent inhibitor of muscle differentiation in chicken and mouse models during embryonic and foetal myogenesis (Bonnet et al., 2010; Delfini and Duprez, 2004; Hirsinger et al., 2001; Mourikis et al., 2012a; Vasyutina et al., 2007; Zalc et al., 2014). Active NOTCH also inhibits muscle differentiation in C2C12 cells (Kopan et al., 1994; Kuroda et al., 1999). NOTCH displays additional functions in postnatal myogenesis, since NOTCH has been shown to be required to maintain the quiescence and to regulate the migratory behaviour of satellite cells (Conboy and Rando, 2002; Conboy et al., 2003, 2005; Baghdadi et al., 2018; Bjornson et al., 2012; Mourikis et al., 2012b). The canonical NOTCH pathway is well described and mediates cell-to-cell communication involving a transmembrane NOTCH receptor and a transmembrane NOTCH ligand. Upon ligand activation, the NOTCH receptor undergoes a proteolytic cleavage to produce the NOTCH intracellular domain (NICD), which translocates to the nucleus and interacts with RBPJ to regulate gene transcription (Andersson et al., 2011; Kopan et al., 1994; Lubman et al., 2007). NICD immediate transcriptome and ChIP-sequencing analyses identified key genes activated downstream of NOTCH, which include the bHLH transcriptional repressors, *Hes* (Hairy and enhancer of split) and *Hey* (Hairy/enhancer-of-split related with YRPW motif) genes (Andersson et al., 2011). In mouse foetal myogenesis, *Heyl* is the main transcriptional response to NICD (Mourikis et al., 2012a). In addition to being primary target genes of NOTCH, HES and HEYL proteins repress the transcription of muscle differentiation genes to maintain muscle cells in a progenitor state in mice (Bröhl et al., 2012; Fukada et al., 2011; Mourikis et al., 2012a; Zalc et al., 2014). Because NOTCH is involved in muscle differentiation, NOTCH function in myoblast fusion is difficult to address. No link has been established between components of the NOTCH intracellular pathway and the fusion gene *myomaker*. Moreover, the location and the molecular regulation of myoblast fusion remain to be understood in vivo.

In this manuscript, we analysed the expression of the key fusion gene *TMEM8C* in addition to that of MYOG+ fusion-competent cells in limb foetal muscles and the consequences of NOTCH inhibition for *TMEM8C* expression and the fusion process during foetal myogenesis using in vitro and in vivo chicken systems.

## Materials and methods

### Chicken embryos

Fertilized chicken eggs from commercial sources (White Leghorn strain, HAAS, Strasbourg, France and JA 57 strain, Morizeau, Dangers, France) were incubated at 38.5°C in a humidified incubator until appropriate stages. Embryos were staged according to the number of days in ovo (E). All experiments on chicken embryos were performed before E14 and consequently are not submitted to a licensing committee, in accordance to the European guidelines and regulations.

### Chemical inhibitor administration in chicken embryos

Decamethonium bromide (DMB) (Sigma, France) solution and control Hank’s solution were prepared as previously described (Esteves de Lima et al., 2016). 100 μl of DMB or control solutions were administered in chicken embryos at E7.5 and E8.5. Embryos were collected at E9.5.

### Grafts of DELTA1/RCAS-expressing cells

Chicken embryonic fibroblasts (CEFs) obtained from E10 chicken embryos were transfected with DELTA1/RCAS using the Calcium Phosphate Transfection Kit (Invitrogen, France). Cell pellets of approximately 50–100 μm in diameter were grafted into limb buds of E4.5 embryos as previously described (Bonnet et al., 2010; Delfini et al., 2000). DELTA1/RCAS-grafted embryos were harvested at E6.5.

### Myoblast cultures

Primary myoblasts were obtained from limbs of E10 chicken embryos and cultured in proliferating high-serum (10%) containing medium or in differentiation low-serum (2%) containing medium, as previously described (Havis et al., 2012). Myoblasts were analysed by immunohistochemistry for PAX7+ muscle progenitors and MF20+ differentiated cells and for muscle gene expression by RT-qPCR. For NOTCH gain-of-function experiments, myoblasts were transfected with DELTA1/RCAS plasmid or Empty/RCAS (control). For NOTCH loss-of-function experiments, myoblasts were treated with DAPT (Sigma) at a concentration of 5 μM for 24 h in low and high serum conditions (proliferation and differentiation assays) versus DMSO (controls). For the fusion assay MYOG+ myoblasts plated at a high density in low serum conditions (Latroche et al., 2017) were treated with 5 μM DAPT at 0 h and 24 h of culture and collected at 48 h.

### Immunohistochemistry

Forelimbs of control or manipulated (DMB, DELTA1/RCAS) chicken embryos were fixed in 4% paraformaldehyde overnight at 4°C and then processed in gelatin/sucrose for 12 μm cryostat sections as previously described (Bourgeois et al., 2015; Esteves de Lima et al., 2014). The monoclonal antibodies, PAX7 and MF20 that recognizes sarcomeric myosin heavy chains, developed by D.A. Fischman and A. Kawakami, respectively, were obtained from the Developmental Studies Hybridoma Bank developed under the auspices of the NICHD and maintained by the University of Iowa, Department of Biology Iowa City, IA 52242. The COL12 and MYOG antibodies were kindly provided by Manuel Koch (Germany) and Christophe Marcelle (France), respectively. Secondary antibodies were conjugated with Alexa-488 or Alexa-555 (Invitrogen). Nuclei were detected with DAPI staining (Sigma).

### In situ hybridization

Chicken forelimbs of control or manipulated (DMB, DELTA1/RCAS) embryos were fixed in Fornoy (60% ethanol, 30% formaldehyde (stock at 37%) and 10% acetic acid) overnight at 4°C and processed for in situ hybridization on wax tissue sections, as previously described (Wang et al., 2010). The digoxigenin-labelled mRNA probes were prepared as previously described: MYOD and DLL1 (Delfini et al., 2000), FGF4 (Edom-Vovard et al., 2001) and MYOG (Bonnet et al., 2010). The TMEM8C probe was obtained by PCR from E9.5 limb tissues using the following primers: Fw: 5’-ACCCTCAGCACTTTGGTCTTT-3’ and Rv: 5’-ACAGGGCACACCCCATACA-3’, cloned into the pCRII-TOPO vector (Invitrogen), linearized with KpnI and synthetized with T7.

### RNA extraction and Real-Time quantitative PCR (RT-qPCR)

Total RNAs were extracted from control limbs, experimental limbs or primary foetal myoblast cultures. 500 ng to 1 μg of RNA was reverse-transcribed using the High-Capacity Retrotranscription kit (Applied Biosystems, France). RT-qPCR was performed using SYBR Green PCR Master Mix (AppliedBiosystems). Primer sequences used for RT-qPCR are listed in Supplementary Table 1. The relative mRNA levels were calculated using the 2^-ΔΔCt method (Livak and Schmittgen, 2001; Schmittgen and Livak, 2008). The ΔCts were obtained from Ct normalized with *GAPDH* and *RPS17* levels in each sample. Each RNA sample was analysed in duplicate.

### Chromatin immunoprecipitation assay

In vitro ChIP assay was performed as previously described (Harada et al., 2018). In vivo ChIP assay was performed as previously described (Havis et al., 2012). Limbs from E9.5 chicken embryos were homogenized using a mechanical disruption device (Lysing Matrix A, Fast Prep MP1, 2×40 s at 6 m/s). 5 μg of the HEYL antibody, kindly provided by So-Ichiro Fukada (Osaka University, Japan), was used to immunoprecipitate 10 μg of sonicated chromatin. ChIP products were analysed by RT-qPCR to amplify the different regions. The primer list is shown in Supplementary Table 1.

### Image capture

After immunohistochemistry or in situ hybridization experiments, images were obtained using a Zeiss apotome epifluorescence microscope, a Leica DMI600B fluorescence microscope or a Leica SP5 confocal system.

### Image analyses (quantification)

The quantification of the number of PAX7+ and MYOG+ cells in control and DAPT foetal myoblast cultures was normalised to the number of DAPI+ cells. The fusion index was performed by quantifying the number of nuclei within myotubes, with a minimum of 2 nuclei, and normalised to the total number of cells (DAPI+). The quantification of the number of nuclei per fibre in vivo was performed by counting the number of nuclei within each fibre. All the described quantifications were performed using the Cell counter plug-in of the free software ImageJ.

### Statistical analyses

Data was analysed using non-parametric two-tailed tests, Mann-Whitney test for unpaired samples or Wilcoxon test for paired samples using Graphpad Prism V6. Results are shown as means ± standard errors or standard errors of the mean indicated in figure legends. The p-values are indicated either with the number on the graphs or with asterisks. Asterisks indicate the different p-values: *p<0.05, **p<0.01, ***p<0.001 and ****p<0.0001.

## Results

### The fusion gene*TMEM8C* and the fusion-competent MYOG+ cells display a regionalised location within foetal skeletal muscles of chicken limbs

The endogenous expression of *TMEM8C* has not been described during development. Given the requirement and sufficiency of *TMEM8C* (*myomaker*) for myoblast fusion (Millay et al., 2013), we used *TMEM8C* transcript location as a readout for muscle fusion in foetal skeletal muscles. In situ hybridization experiments to longitudinal limb sections of E8.5 chicken embryos showed *TMEM8C* expression (Fig. 1A,C) in limb foetal muscles visualised with *MYOD* (Fig. 1B,D) or myosins (Fig. 1E). *TMEM8C* displayed a higher expression within the middle of the muscles versus the muscle ends, labelled with *FGF4* transcripts (Fig. 1I), which was identified as being regionalized in muscle regions close to tendons (Edom-Vovard et al., 2001, 2002). Comparison of *TMEM8C* expression with that of *FGF4* indicated a complementary expression pattern; i.e. a central muscle location for *TMEM8C* and muscle extremities for *FGF4* (Fig. 1H,I). *TMEM8C* expression was observed inside and outside myosin+ myotubes (Fig. 1F,G arrowheads and arrows, respectively). Because of the strong expression of *TMEM8C* in central muscle regions in limb muscles (Fig. 1), we investigated the location of the differentiated and fusion-competent MYOG+ myoblasts within muscles. Immunohistochemistry on transverse limb sections showed fewer MYOG+ nuclei in muscle regions close to tendon attachments, which were visualized with type XII collagen, compared to the central muscle regions where MYOG+ cells were more abundant (Fig. 2A,B). High magnification observations of a ventral and posterior limb muscle, the FCU (*Flexor Carpi Ulnaris*), confirmed the high density of MYOG+ cells in central muscle regions compared to muscle ends (Fig. 2C), while PAX7+ muscle progenitors displayed a more homogenous distribution in muscle (Fig. 2D). Analysis of longitudinal muscle sections also showed a lower density of MYOG+ nuclei in muscle regions close to tendon attachments, compared to central regions, while PAX7+ muscle progenitors were more homogenously distributed in limb muscles (Fig. S1).

**Figure 1.**
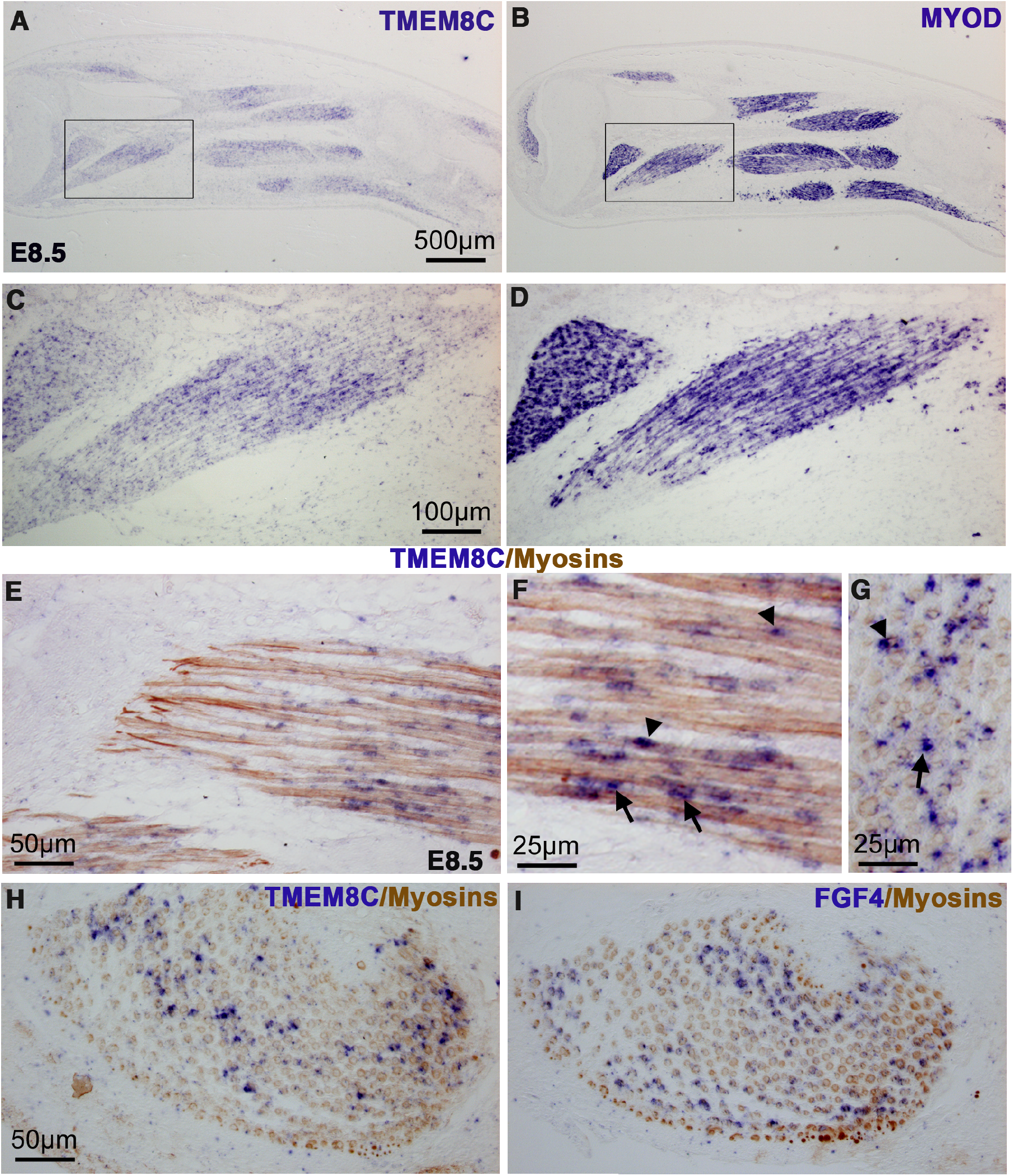
*TMEM8C* expression in foetal skeletal muscles of chicken limbs. (A-D) In situ hybridization to adjacent and longitudinal limb sections of E8.5 chicken embryos with the TMEM8C (A,C) and MYOD (B,D) probes. (C,D) are high magnifications of the squared regions in (A,B), respectively. *TMEM8C* displays a central and more restricted expression compared to that of *MYOD* in muscles. (E-G) Longitudinal (E,F) and transverse (G) limb muscle sections were hybridized with the TMEM8C probe (blue) followed by immunohistochemistry with the MF20 antibody to visualize myosins (brown). (F) is a higher magnification of (E). *TMEM8C* transcripts (blue) are observed inside (arrowheads) and outside (arrows) myotubes. (H,I) In situ hybridization to adjacent and transverse limb sections of E8.5 chicken embryos with the TMEM8C (H) and FGF4 (I) probes. Central *TMEM8C* expression is complementary to that of *FGF4* expression at muscle extremities.

**Figure 2.**
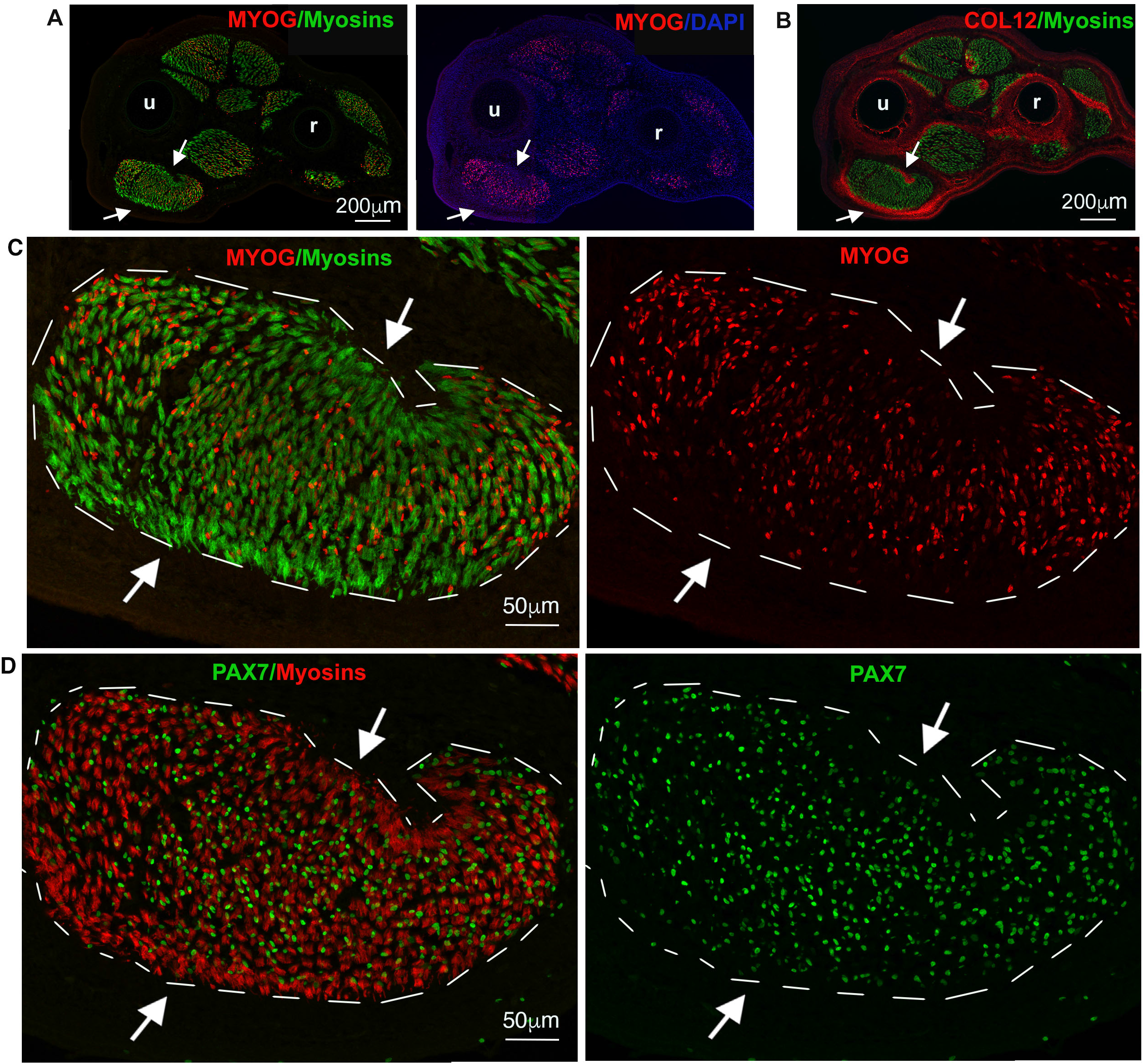
Regionalisation of PAX7+ and MYOG+ nuclei in foetal muscles of chicken limbs. (A,B) Immunohistochemistry to adjacent and transverse limb sections of E9 chicken embryos with antibodies for MYOG (A), type XII collagen (COL12), which labels tendons (B) and myosins (MF20 antibody) (A,B). Arrows in (A,B) point to tendons of the FCU, a muscle located in ventral and posterior limb regions. (C) is a high magnification of the FCU muscle shown in (A), labelled with MYOG and MF20 antibodies. (C) Dashed lines delineate the myosin+ area of the FCU to highlight MYOG+ cell regionalisation within muscles. Arrows in (C) indicate muscle regions, close to tendons, which display a low density of MYOG+ cells. (D) Adjacent sections to (C) show FCU muscle immunolabelled with PAX7 and MF20 antibodies. (D) Dashed lines delineate the myosin+ area of the FCU to highlight the PAX7+ cells within muscles. u, ulna; r, radius.

The similar location of the *TMEM8C* transcripts (Fig. 1) and fusion-competent MYOG+ cells (Fig. 2) suggests than myoblast fusion occurs preferentially in central muscle regions of limb foetal muscles.

### NOTCH loss-of-function following embryo immobilisation increases muscle fusion in foetal limb muscles

Chicken embryo immobilisation leads to a NOTCH loss-of-function phenotype in limb muscles, phenotype that can be rescued with DELTA1-activated NOTCH in immobilisation conditions (Esteves de Lima et al., 2016). Immobilisation induces a drastic decrease in the expression levels of the NOTCH ligand *JAG2* in differentiated muscle cells and a decrease in the expression of transcriptional readout of NOTCH activity, *HEYL* in limb muscles. This results in a shift towards muscle differentiation associated with a decrease in the number of PAX7+ progenitors (Esteves de Lima et al., 2016). In the same immobilisation conditions, following decamethonium bromide (DMB) treatment, we observed an increase in the expression of the *TMEM8C* fusion gene in limb foetal muscles compared to controls (Fig. 3A-D). The increase of *TMEM8C* mRNA levels in paralysed limb muscles was confirmed by RT-qPCR (Fig. 3E). As expected, the *PAX7* expression levels were decreased, while those of *MYOG* increased (Fig. 3E). The quantification of the number of myonuclei per fibre showed a significant increase in muscles of immobilised embryos compared to mobile embryos (Fig. 3F-H), indicating an increased incorporation of nuclei in myofibres of paralyzed muscles. Altogether, these experiments show that NOTCH loss-of-function following embryo immobilisation increases muscle fusion in addition to differentiation in limb foetal muscles of chicken embryos.

**Figure 3.**
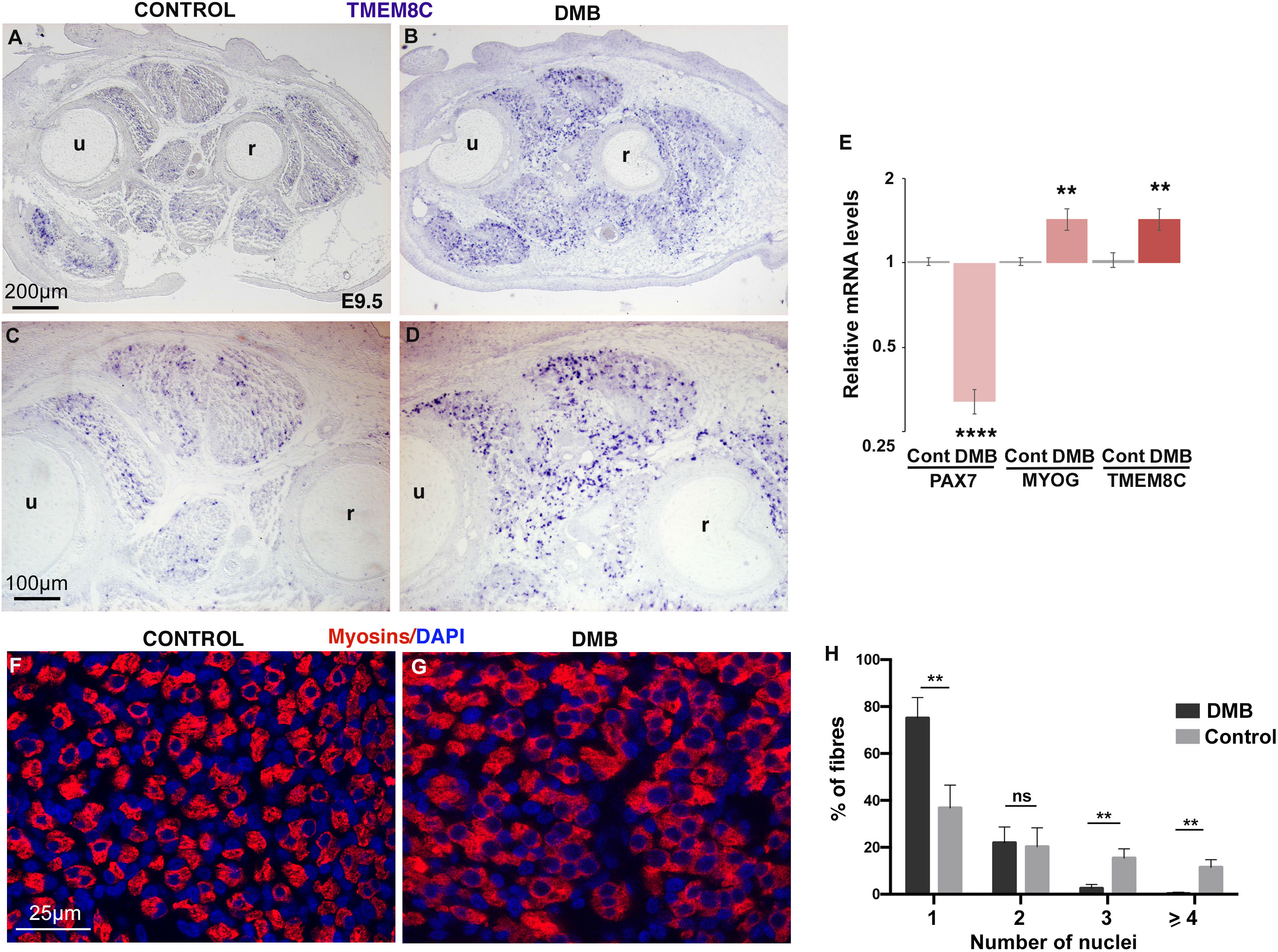
NOTCH loss-of-function following immobilisation of chicken embryos increases *TMEM8C* expression and fusion in limb muscles. DMB was administrated in chicken embryos at E7.5 and E8.5 to induce immobilisation for 48 h. Limbs of immobilised embryos for 48 h (E9.5) were either processed for in situ hybridisation (A-D), RT-qPCR analyses (E) or immunohistochemistry (F,G). (A-D) Forelimb transverse sections of control (n=4) (A,C) and DMB-treated (n=4) (B,D) embryos were hybridised with TMEM8C probe. (C,D) are high magnifications of dorsal limb muscles of (A,B), respectively. (E) RT-qPCR analyses of the mRNA levels for *PAX7, MYOG* and *TMEM8C* genes in control and paralysed limbs. Graph shows means ± standard error of the mean of gene expression in 10 control forelimbs and 10 paralysed forelimbs. The relative mRNA levels were calculated using the 2^^−ΔΔCt^ method. For each gene, the mRNA levels of control limbs were normalised to 1. (F,G) Transverse sections of limb muscles from mobile (n=3) (F) and immobilised (n=3) (G) embryos were immunostained with MF20 and DAPI to visualise nuclei in myofibres. (H) Quantification of the number of nuclei in myofibres observed in mobile and immobilised embryos. Error bars represent standard deviation. \*\**p*<0.01; \*\*\*\**p*<0.0001. u, ulna; r, radius.

### Validation of an in vitro system of chicken foetal myoblast cultures that mimics in vivo myogenesis

The processes of differentiation and fusion are not easy to dissociate in vivo during skeletal muscle development, therefore we developed an in vitro system of chicken foetal myoblast cultures to dissect these two processes. Foetal myoblasts were isolated from limbs of E10 chicken embryos and plated at low density with high serum-containing medium defining proliferation conditions. Muscle cell cultures in proliferation conditions contained PAX7+ progenitor cells and myosin+ differentiated cells (Fig. S2A,B). Confluent myoblasts pushed to differentiation and fusion with low serum-containing medium formed multinucleated myosin+ cells and kept a pool of PAX7+ reserve cells (Fig. S2C,D). The analysis of muscle gene expression showed that the expression of the progenitor markers *PAX7* and *MYF5* was decreased, while that of *MYHC* was increased in differentiation conditions compared to proliferating conditions (Fig. S2E). In order to assess NOTCH activity during muscle differentiation in myoblast cultures, we analysed the expression levels of the NOTCH transcriptional readout *HEYL*. Consistently with the in vivo situation, *HEYL* expression was decreased in differentiated muscle cell cultures compared to myoblasts in proliferation culture conditions (Fig. S2E).

We then asked if the manipulation of NOTCH activity would have similar outcomes for myogenesis in vitro and in vivo. To test this, we performed NOTCH loss- and gain-of-function experiments in myoblasts cultured in proliferation conditions and compared the phenotypes obtained with those observed in vivo in chicken limbs. We first performed NOTCH loss-of-function experiments in myoblasts using the gamma-secretase inhibitor DAPT. DAPT prevents the proteolytic cleavage of NICD, which is a necessary step for the NOTCH signalling response, as NICD translocates to the nucleus to regulate gene transcription. NOTCH inhibition with DAPT treatment increased myoblast differentiation and increased the expression levels of the differentiation markers *MYOD, MYOG* and *MYHC,* while decreasing those of *PAX7* and *MYF5* in proliferating myoblasts (Fig. 4A,B,D). This result is consistent with the myogenic phenotype (increase of *MYOD* and *MYOG* expression and decrease in the number of PAX7+ cells) generated by NOTCH-loss-of-function observed in limb foetal muscles after immobilisation of chicken embryos (Esteves de Lima et al, 2016). Conversely, we forced NOTCH activity by overexpressing DELTA1 using the RCAS retroviral system in myoblasts and in chicken limbs. DELTA1-activated NOTCH led to the mirror phenotype for myogenesis to that of DAPT experiments in myoblasts. DELTA1-activated NOTCH increased the number of PAX7+ cells in myoblasts cultured in proliferation conditions (Fig. S3A,B) and in chicken limbs after two days of DELTA1/RCAS exposure (Fig. S3C-H) compared to controls. The expression levels of *MYOD, MYOG* and *MYHC* mRNAs were decreased, while those of *PAX7* and *MYF5* were increased in DELTA1-activated NOTCH myoblast cultures (Fig. 4A,C,E). This was consistent with the decreased expression of *MYOD* and *MYOG* transcripts and myosin protein previously described in DELTA1-activated NOTCH limbs of chicken embryos (Bonnet et al., 2010; Delfini et al., 2000). Upon altered NOTCH activity, the NOTCH target gene *HEYL* was accordingly mis-regulated in myoblasts, but not those of BMP signalling *ID1* and *ID2* (Fig. 4D,E), previously shown to interact with NOTCH in other systems (Guo and Wang, 2009).

**Figure 4.**
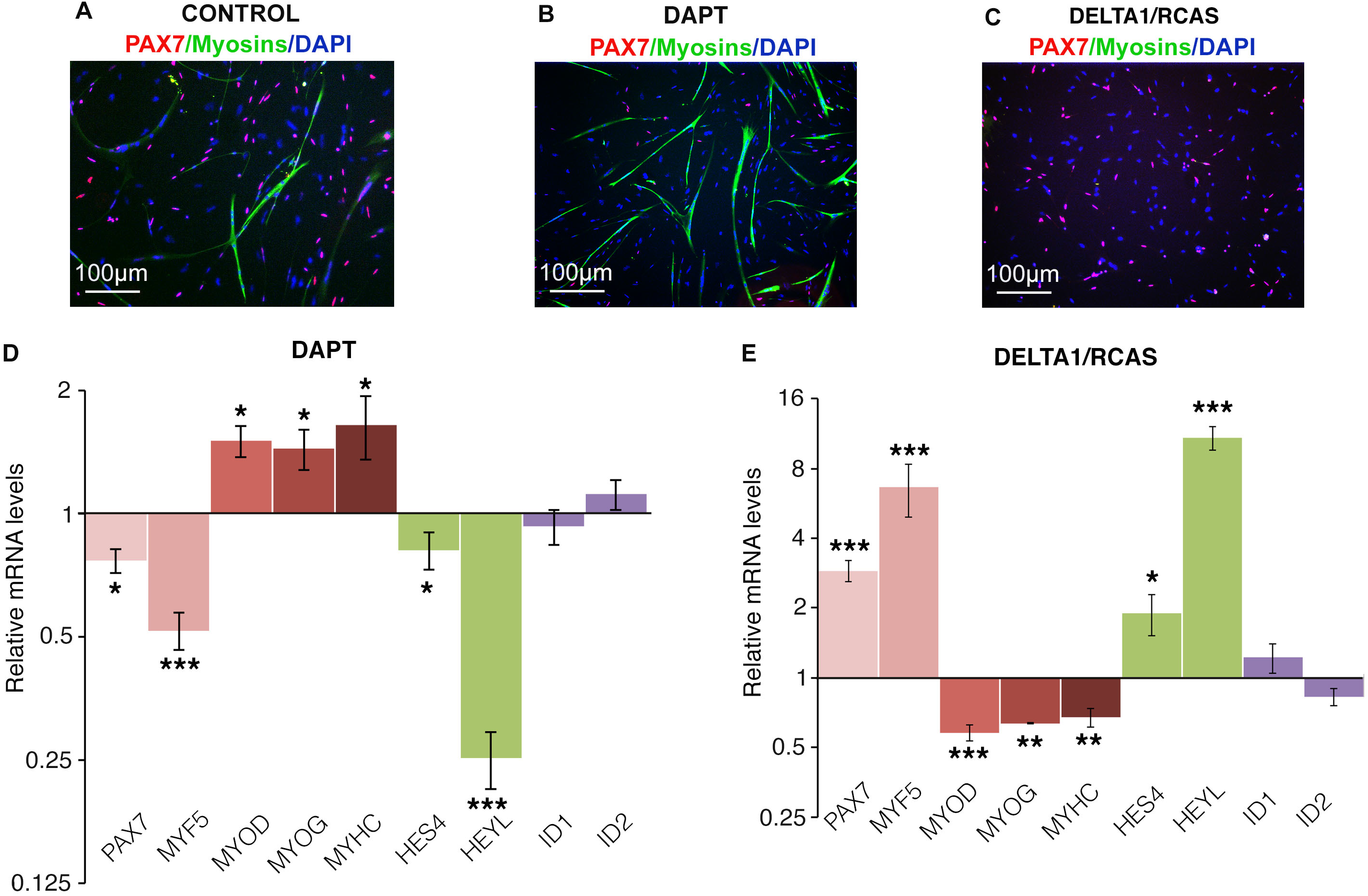
NOTCH inhibition and activation experiments lead to opposite effects in myoblasts cultured in proliferation conditions. (A,B,C) Representative fields showing PAX7+ and myosin+ cells in chicken foetal myoblasts, transfected with Empty/RCAS (A), treated with DAPT (B) or transfected with DELTA1/RCAS (C), and cultured in proliferation conditions. (D,E) RT-qPCR analyses of the expression levels for muscle markers (*PAX7, MYF5, MYOD*, *MYOG, MYHC*), NOTCH target genes (*HES4, HEYL*) and BMP target genes (*ID1, ID2*) in DAPT-treated (D) and DELTA1/RCAS-infected (E) myoblasts cultured in proliferation conditions (n=7). The relative mRNA levels were calculated using the 2^^−ΔΔCt^ method. For each gene, the mRNA levels of RCAS-infected myoblasts (control) were normalised to 1. Error bars indicate standard error of the mean. *p<0.05; ** p<0.01; *** p<0.001.

These results show that cultures of chicken foetal myoblasts mimic in vivo myogenesis and that NOTCH loss- and gain-of-function lead to similar outcomes for muscle progenitor number and gene expression in vivo and in vitro.

### NOTCH loss-of-function experiments in myoblasts increased myoblast fusion in addition to differentiation

In order to assess the effect of NOTCH inhibition on differentiation and fusion, myoblast cultures were led to differentiate in low-serum conditions with DMSO or DAPT for 24 h (Fig. 5). DAPT-mediated NOTCH inhibition favoured the appearance of myotubes and a decrease in the number of PAX7+ cells as compared to controls (Fig. 5A,B). Consistently, the expression of the muscle differentiation marker *MYOG* was increased, while that of *PAX7* was decreased, in DAPT-treated cultures compared to control cultures (Fig. 5C). In addition, *TMEM8C* expression was also increased in DAPT-treated cultures (Fig. 5C). Consistent with the increase of *TMEM8C* mRNA levels (Fig. 5C), the myotubes were bigger and the fusion index was significantly increased in DAPT-treated myoblasts cultured in differentiation conditions compared to controls (Fig. 5D,E). These results show that NOTCH inhibition increases the differentiation and fusion processes in foetal myoblasts cultured in differentiation conditions.

**Figure 5.**
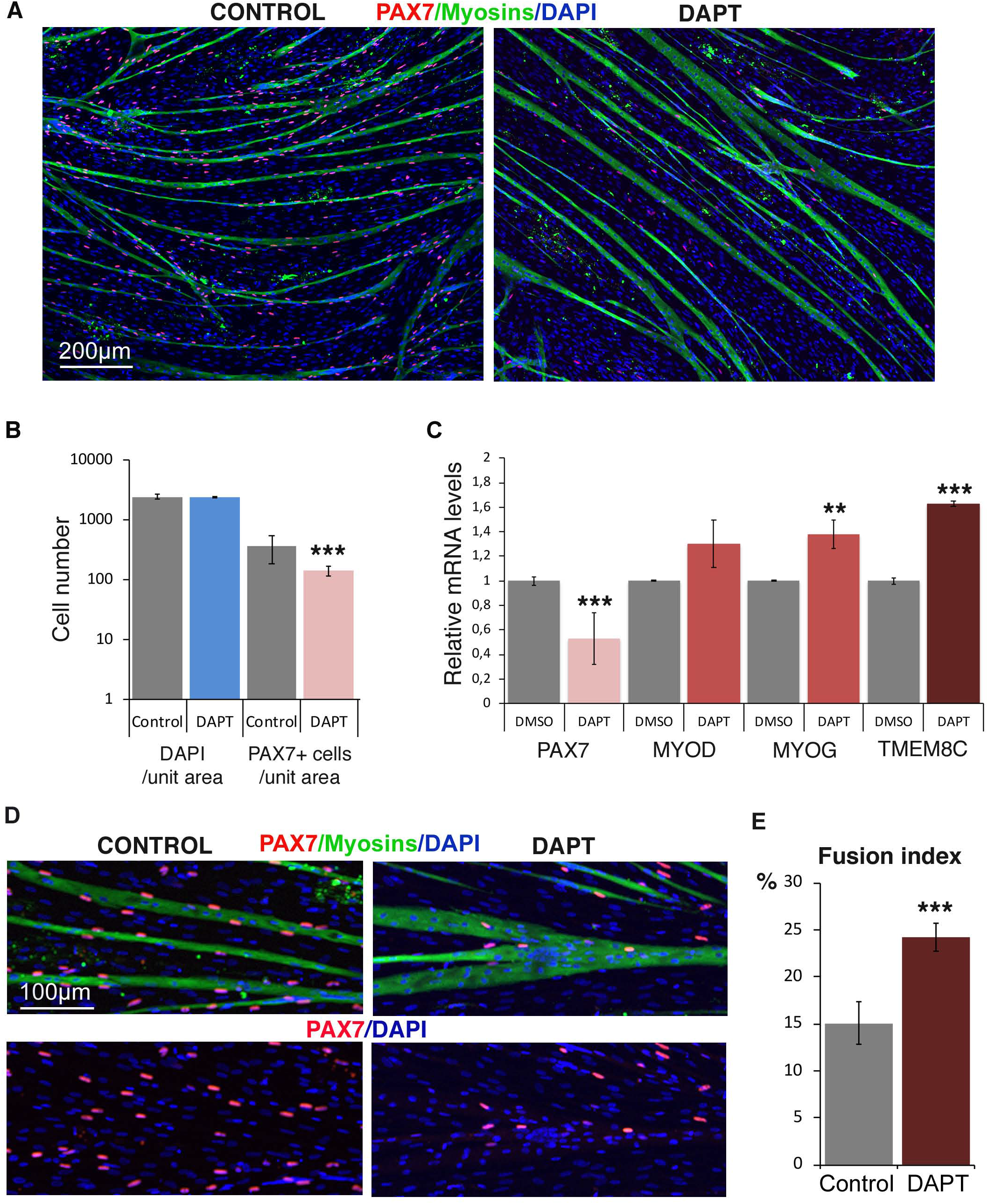
Inhibition of NOTCH activity promotes differentiation and fusion in foetal myoblasts cultured in differentiation conditions. (A) Representative fields of myotubes of chicken myoblast cultures in differentiation conditions treated with DMSO (control) or DAPT, labelled with PAX7 (muscle progenitors) and MF20 (myosins) antibodies combined with DAPI staining (nuclei). (B) Quantification of the number of nuclei and PAX7+ cells per unit area. Graph shows means ± standard deviation of 5 biological samples. (C) RT-qPCR analyses of the expression levels for muscle markers, *PAX7, MYOD*, *MYOG* and *TMEM8C* in control and DAPT-treated myoblasts cultured in differentiation conditions. Graph shows means ± standard deviation of 9 control and 7 DAPT-treated myoblast cultures. The relative mRNA levels were calculated using the 2^^−ΔΔCt^ method. For each gene, the mRNA levels of control myoblasts were normalised to 1. (D) High magnification showing myotubes labelled with MF20 antibody and PAX7+ progenitors combined with DAPI staining of chicken muscle cells cultured in differentiation conditions treated with DMSO (control) or with DAPT. (E) Fusion index of chicken muscle cells cultured in differentiation conditions treated with DMSO (control) or with DAPT (n=4). Error bars indicate standard deviations. ** p<0.01; *** p<0.001.

### NOTCH loss-of-function increased fusion in differentiated myoblasts

In the previous NOTCH inhibition experiments (Fig. 5), we could not dissociate muscle cell differentiation from fusion in myoblasts cultured in differentiation conditions. In order to uncouple the initial step of differentiation from fusion, we used a two-step protocol where proliferating myoblasts were first pushed to differentiation without fusion and then allowed to fuse in a second step (Fig. 6A). Based on previously described protocols (Girardi et al., 2019; Latroche et al., 2017), we induced differentiation with low serum medium in myoblasts cultured in a sub-confluent manner to avoid cell-cell contact, with the idea that proliferating myoblasts will enter the differentiation program without undergoing fusion (Fig. 6A). As expected, after this procedure, myoblasts became MYOG+ and lost PAX7 expression (Fig. 6B-D). MYOG+ differentiated cells were then plated at high density and treated with either DAPT or DMSO for 48 h to assess the effect of NOTCH inhibition on the fusion process of already-differentiated myoblasts. DAPT-mediated NOTCH inhibition increased myoblast fusion and myotube formation in differentiated myoblasts compared to controls (Fig. 6E,F). The fusion index was increased 2-fold in the context of NOTCH inhibition compared to control cultures (Fig. 6G). Taken together, these results show that NOTCH inhibition promotes fusion of differentiated myoblasts in vitro.

**Figure 6.**
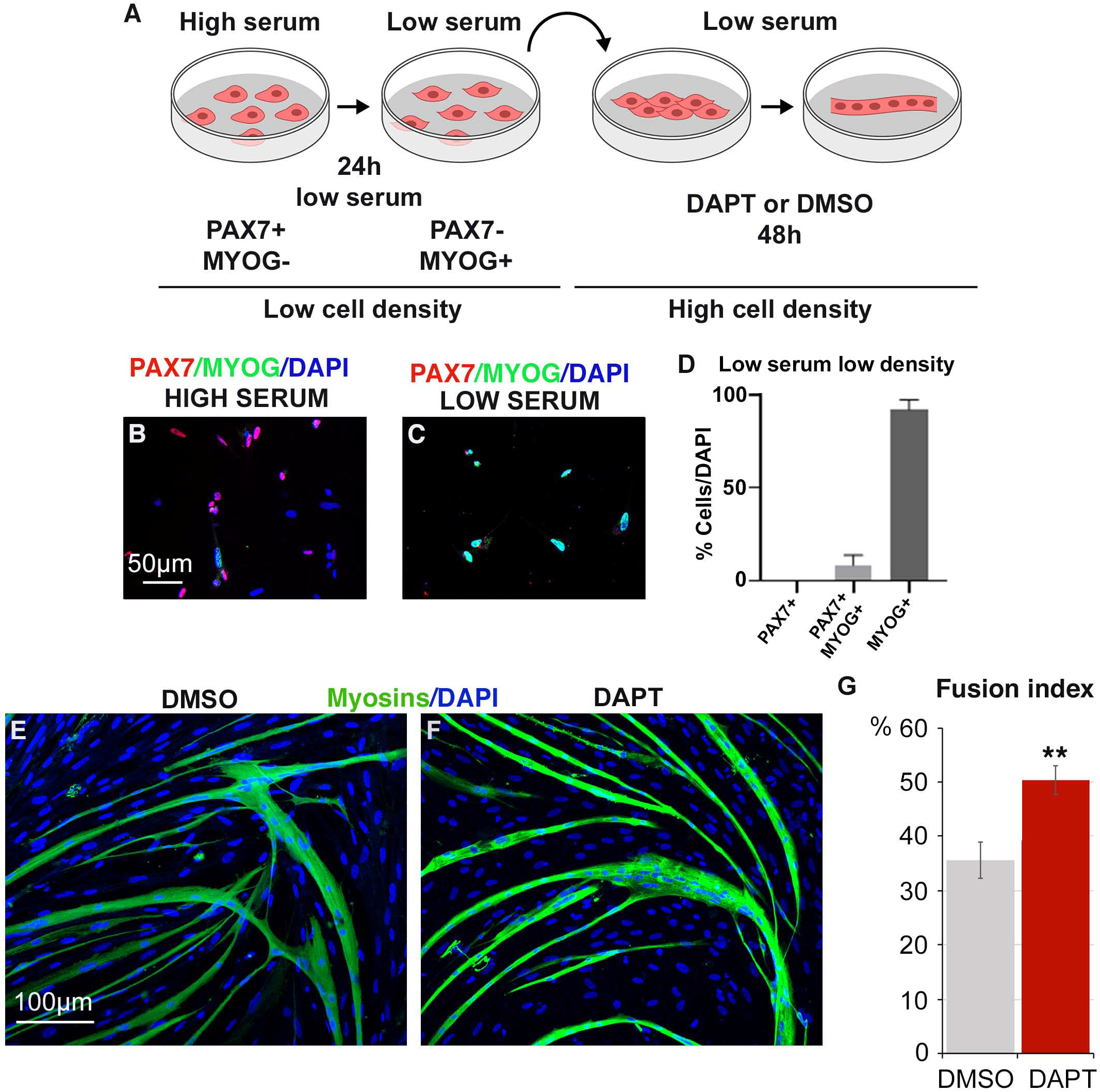
Inhibition of NOTCH activity promotes fusion in differentiated foetal myoblasts. (A) Scheme of the experimental design to promote myoblast differentiation without fusion. (B,C) Myoblasts cultured at low density with proliferation medium (B) and differentiation medium (C), labelled with PAX7 (red) and MYOG (green) antibodies. (D) Quantification of PAX7+ and MYOG+ cells in myoblasts cultured at low cell density and with low serum. (E,F) MYOG+ differentiated myoblasts plated at high density were treated with DMSO (E) or DAPT (F) for 48 h and immunostained with MF20 to label myosins. (G) Fusion index of differentiated myoblasts plated at high density treated with DMSO (control) or with DAPT (n=6). Error bars indicate standard error of the mean. ** p<0.01.

### NOTCH loss-of-function in limbs and myoblast cultures released HEYL repressor recruitment to*TMEM8C* regulatory regions

In order to define the molecular mechanism through which NOTCH inhibition promoted myoblast fusion, we analysed the recruitment of the NOTCH target gene HEYL to the *TMEM8C* regulatory regions. Because the HEY/HES are bHLH transcriptional repressors known to inhibit myogenesis via the direct binding to E-boxes-containing regions of the *MYOD* enhancer (Zalc et al., 2014), we tested if HEYL could bind to the 3 E-boxes-containing regions in regulatory regions of the *TMEM8C* fusion gene (Fig. 7A). In order to test this hypothesis, we performed chromatin immunoprecipitation (ChIP) experiments in two NOTCH-inhibition conditions: (1) cultures of differentiated myoblasts treated with DAPT (two-step protocol) and (2) limb muscles of immobilised chicken embryos. We found that in both NOTCH-inhibition conditions, the recruitment of the transcriptional repressor HEYL to the 3 E-box-containing regions located upstream of *TMEM8C* was decreased compared to control conditions (Fig. 7B,C). This result is consistent with the increase of *TMEM8C* expression in DAPT-treated myoblasts (Fig. 5C) and in limb foetal muscles after embryo immobilization (Fig. 3A-E). The decrease of HEYL recruitment to the *TMEM8C* promoter in the absence of NOTCH activity in vitro and in vivo provides us with a potential mechanism for the fusion-promoting effect of NOTCH inhibition (Fig. 7D).

**Figure 7.**
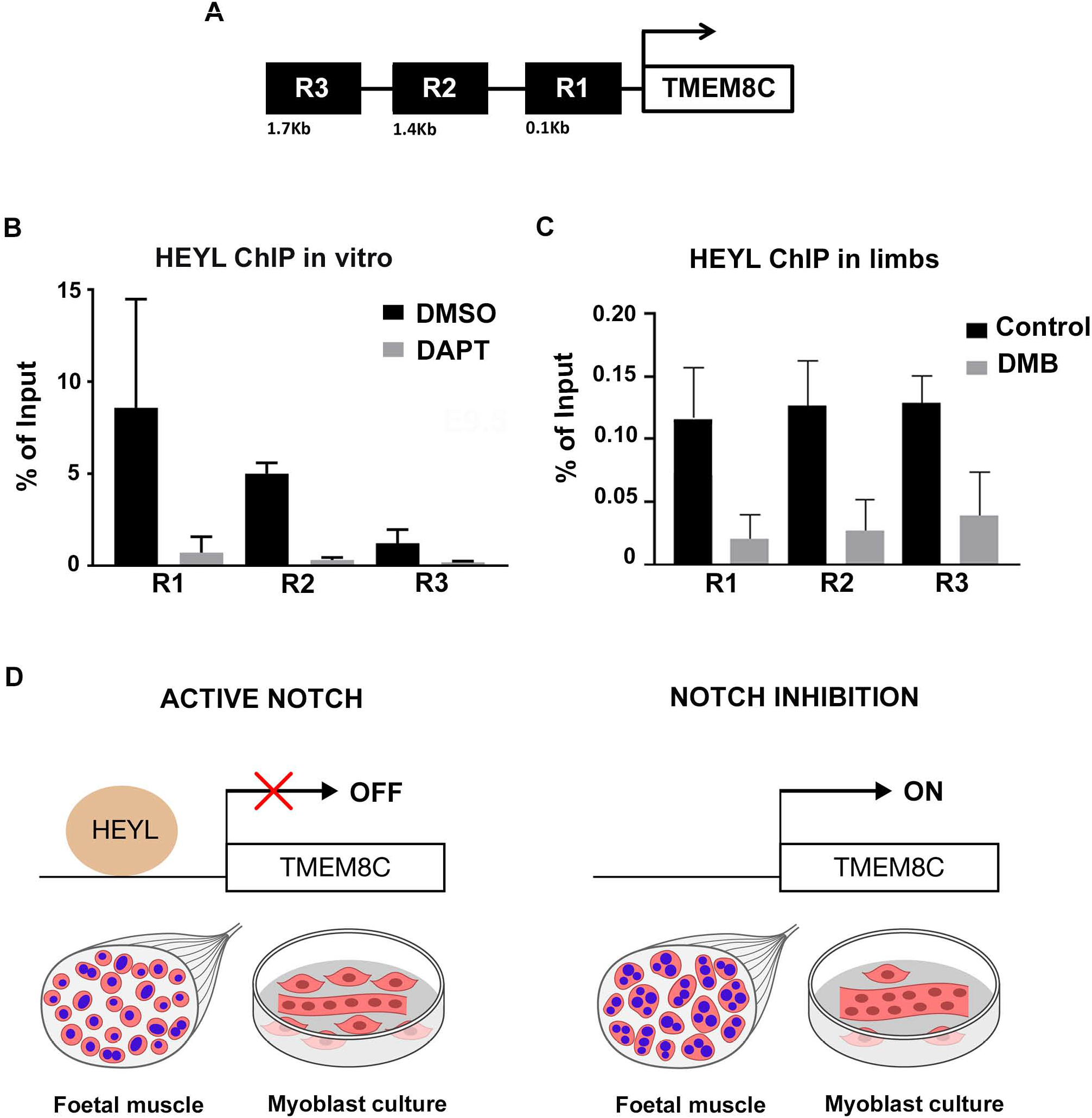
Recruitment of HEYL repressor is released from *TMEM8C* regulatory regions in limb muscles of immobilized chicken embryos and DAPT-treated differentiated myoblast cultures. (A) Scheme of the 3 E-box-containing regions upstream the transcription starting site of the *TMEM8C* gene, tested by ChIP-qPCR, R1 (+36 bp – −272 bp), R2 (−1.2 kb – −1.5 kb) and R3 (−1.5 kb – −1.7 bp). (B,C) ChIP assays were performed on differentiated myoblast cultures treated with DAPT or DMSO (n=4) (B) and on chicken limb muscles of mobile and immobile embryos mimicking a NOTCH loss-of-function phenotype (n=3) (C), with an antibody against HEYL to analyse HEYL recruitment to *TMEM8C* regulatory regions. HEYL was recruited to the 3 E-box-containing regions of *TMEM8C* gene in control conditions, while the HEYL recruitment was released upon NOTCH inhibition in vivo and in vitro. Error bars indicate standard deviations. (D) Schematic representation of the recruitment of the transcriptional repressor HEYL to the *TMEM8C* regulatory regions in control and NOTCH inhibition conditions.

## Discussion

In the present study we identified a regionalised location for the *TMEM8C* fusion gene and fusion-competent MYOG+ cells in limb foetal muscles of chicken embryos and established a molecular link between the NOTCH pathway and the fusion gene *TMEM8C* that could be the basis for the fusion-promoting effect of NOTCH inhibition.

Whether the fusion process occurs at specific places within the skeletal muscles is not known. We identified a preferential central location for *TMEM8C* transcripts and fusion-competent MYOG+ cells in skeletal muscles during foetal myogenesis in chicken (Fig. 1,2). The similar location of MYOG+ cells and *TMEM8C* transcripts is fully consistent with the transcriptional regulation of *TMEM8C* gene by MYOG (Luo et al., 2015; Millay et al., 2014; Ganassi et al., 2018). This regionalized expression suggests that fusion would preferentially occur in the middle of muscle during chicken foetal development. In apparent contradiction, based on BrdU incorporation experiments, myoblast fusion has been suggested to occur preferentially at muscle ends in rat and mouse muscles during foetal and perinatal growth (Gu et al., 2016; Kitiyakara and Angevine, 1963; Zhang and McLennan, 1995). However, the absence of clear definition of muscle extremities can contribute to different interpretations of the fusion location. Moreover, the models proposed in these studies did not take into account the possible migration of myonuclei within the fibre, which can bias the analysis based on the presence versus absence of BrdU+ myonuclei. Furthermore, it has been shown a preferential location of proliferating PAX7+ cells in muscle regions close to tendons, indicating that the muscle end environment favours a proliferative state of the myoblasts rather than fusion (Esteves de Lima et al., 2014).

Because NOTCH inhibition is a potent trigger of muscle differentiation, the role of NOTCH in muscle fusion has been neglected. We now show that NOTCH inhibition increased myoblast fusion in addition to differentiation in two systems, in foetal myoblast cultures and limb foetal muscles after embryo immobilisation. This is consistent with the myotube hypertrophy observed in mouse C2.7 myoblasts upon NOTCH inhibition (Kitzmann et al., 2006). Moreover, most of the molecules identified as being involved in myoblast fusion can be linked to NOTCH signalling. The calcium-activated transcription factor NFATC2, recognized to control myoblast fusion after the initial formation of myotubes (Horsley et al., 2003), has the ability to suppress NOTCH transactivation and the expression of *Hey* genes (Zanotti et al., 2013). Shisa2, an endoplasmic reticulum (ER) localized protein, which regulates the fusion of mouse myoblasts via Rac1/Cdc42-mediated cytoskeletal F-actin remodelling, is repressed by NOTCH signalling (Liu et al., 2018). SRF was recently identified as a regulator of satellite cell fusion via the maintenance of actin cytoskeleton architecture (Randrianarison-Huetz et al., 2018). The NOTCH target gene Herp1 physically interacts with SRF to interfere with SRF transcriptional activity (Doi et al., 2005). Recently, it has been reported that TGFβ inhibition promotes muscle cell fusion in chicken embryos and adult mouse muscles by modulating actin dynamics (Girardi et al., 2019; Sieiro et al., 2019). Interestingly, crosstalks have been identified between TGFβ and NOTCH intracellular signalling pathways, which lead to functional synergism for both pathways. TGFβ cooperates with NOTCH to induce *Hes1, Hey1* and *Jag1* expression in a Smad3-dependent manner through a Smad3–NICD interaction in different systems (Blokzijl, 2003; Zavadil et al., 2004). Given this positive interaction, we cannot exclude that TGFβ inhibition interferes with NOTCH signalling or vice versa that NOTCH inhibition interferes with TGFβ signalling. NOTCH decay has been also involved in the fusion process of fusion-competent myoblasts into myotubes in adult Drosophila (Gildor et al., 2012), suggesting a generic involvement of NOTCH inhibition in myoblast fusion in invertebrates and vertebrates.

In addition to showing a fusion-promoting effect of NOTCH inhibition in vertebrates, we established a molecular link between NOTCH and the fusion gene *TMEM8C (myomaker).* In control conditions, the recruitment of the HEYL repressor to the E-box-containing regions of the *TMEM8C* gene could be the basis for the absence of fusion ability of muscle progenitors displaying active NOTCH (Figure 7D). In the context of NOTCH inhibition, the expression of the NOTCH target gene *HEYL* is decreased and consequently the HEYL recruitment to *TMEM8C* regulatory regions is lost, which results in the release of the inhibition of *TMEM8C* expression by the transcriptional repressor HEYL (Figure 7D). The increase of *TMEM8C* expression is likely to be the molecular mechanism underlying the increased fusion observed in NOTCH inhibition conditions. This also indicates that during normal development, when NOTCH activity is decreased in muscle cells, muscle fusion is promoted in addition to differentiation. The differentiation factors MYOD/MYOG also bind to the E-box domain located close to the transcription starting site of the *myomaker* gene to positively regulate *myomaker* expression in mouse (*Mymk*), chicken (*TMEM8C)* and zebrafish (*mymk*) (Luo et al., 2015; Millay et al., 2014; Ganassi et al., 2018). One attractive mechanism could be a competition between HEYL transcriptional repressor and MYOG transcriptional activator for the occupancy of the *TMEM8C* regulatory regions. In the context of active NOTCH, MYOG is not present and HEYL represses *TMEM8C* transcription, while in the context of NOTCH inhibition, HEYL is decreased and MYOG is present and activates *TMEM8C* transcription. This provides us with a possible new mechanism for tuning myoblast differentiation and fusion in myogenesis during development.

## Acknowledgements

We thank lab members, Matthew Borok, Philippos Mourikis, Sonya Nassari and Valentina Taglietti for critical reading of the MS. We thank So-Ichiro Fukada, Manuel Koch and Christophe Marcelle for reagents. We thank Sophie Gournet for illustrations. The work was supported by the AFM MyoSig N°15761 and AFM Myoconnect N°16752, ANR BMP-MyoMass and FRM DEQ20140329500 and CNRS, UPMC, INSERM. JEdL was part of the MyoGrad International Research Training Group for Myology and was supported by the DFG and the AFM.

---

**Figure S1.**
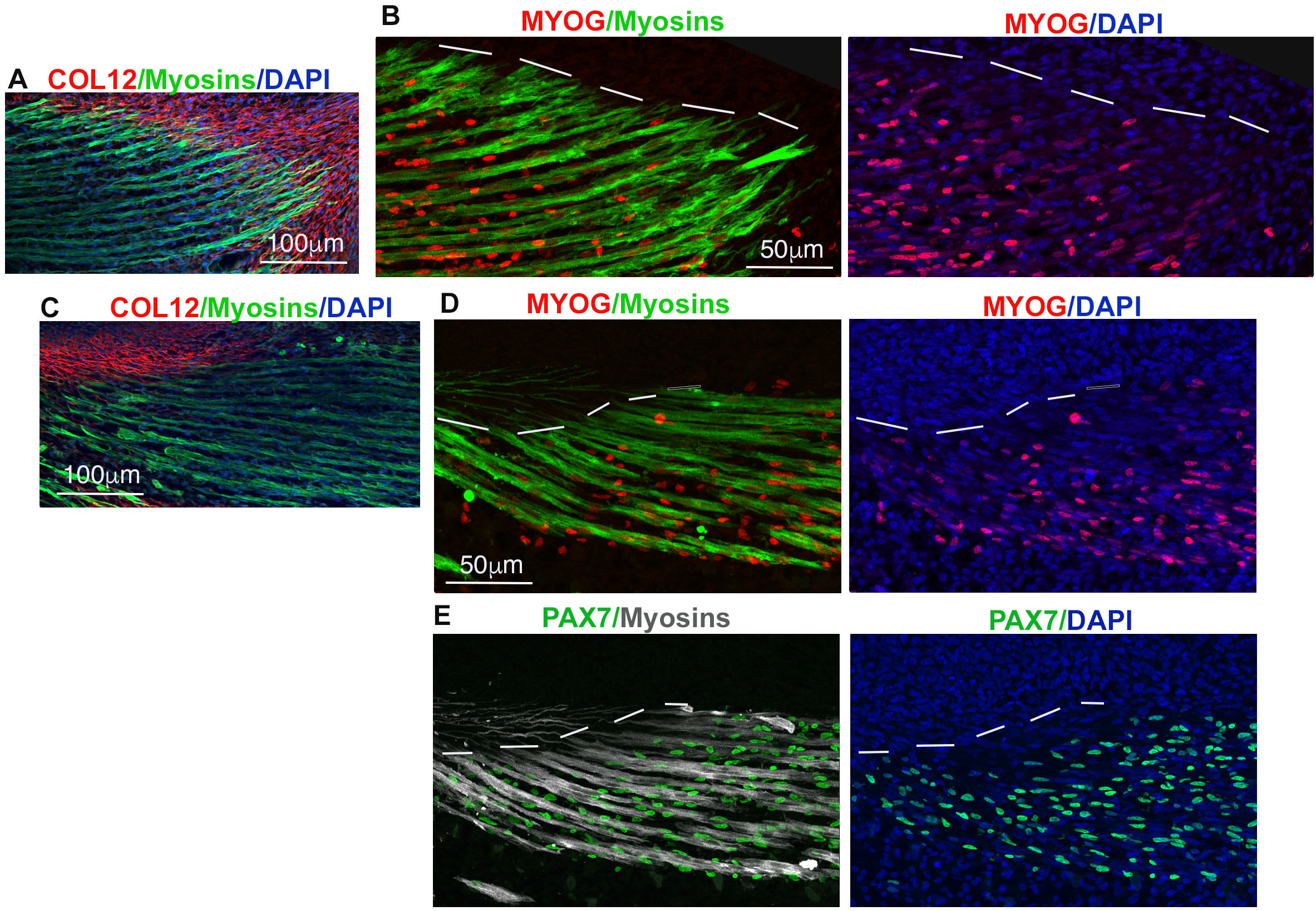
MYOG+ nuclei are regionalised in foetal muscles of chicken limbs. (A,B) Immunohistochemistry to adjacent and longitudinal limb sections of E9 chicken embryos with type XII collagen (COL12 in red) expressed in tendons (A) and MYOG (red) and MF20 (recognising myosins in green) (B) antibodies. (C-E) Immunohistochemistry to adjacent and longitudinal limb sections of E9 chicken embryos with antibodies recognizing type XII collagen (COL12 in red) (C), MYOG (red) and myosins (green) (D), and PAX7 (green) and myosins (grey) (E). Dashed lines delineate the border between muscle and tendon.

**Figure S2.**
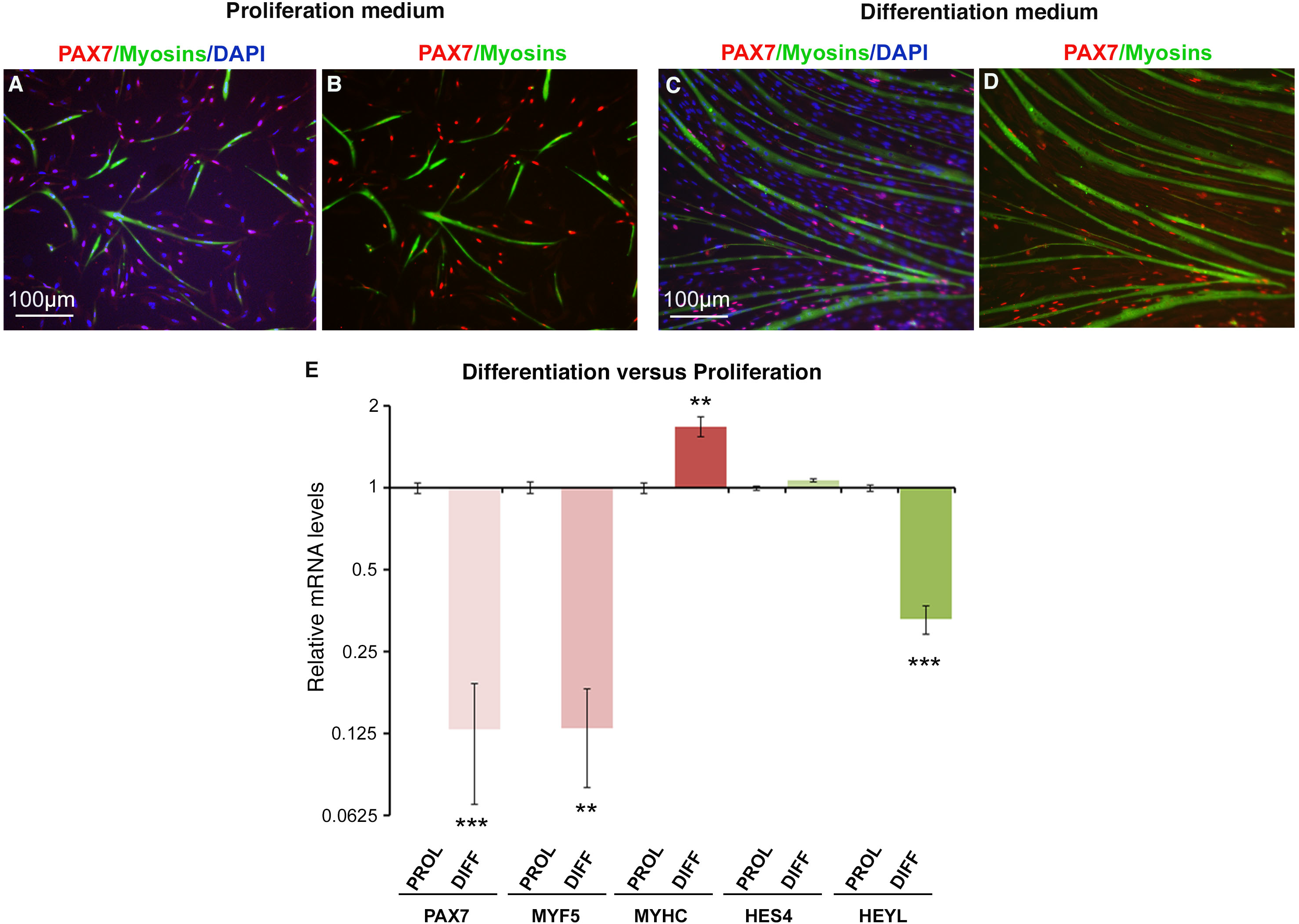
NOTCH activity is downregulated in chicken myoblast cultures during differentiation. (A-D) Representative fields of proliferating (A,B) and differentiating (C,D) chicken myoblasts labelled with PAX7 and MF20 antibodies. (E) RT-qPCR analyses of the mRNA expression levels for the muscle markers, *PAX7, MYF5, MYHC* and the NOTCH target genes, *HES4, HEYL*, in proliferation and differentiation culture conditions. The mRNA levels of the genes of myoblasts in proliferation conditions were normalised to 1 and the relative mRNA levels of the genes in differentiation conditions was calculated versus the expression in proliferating conditions (n=10). Error bars indicate standard error of the mean. ** p<0.01; *** p<0.001.

**Figure S3.**
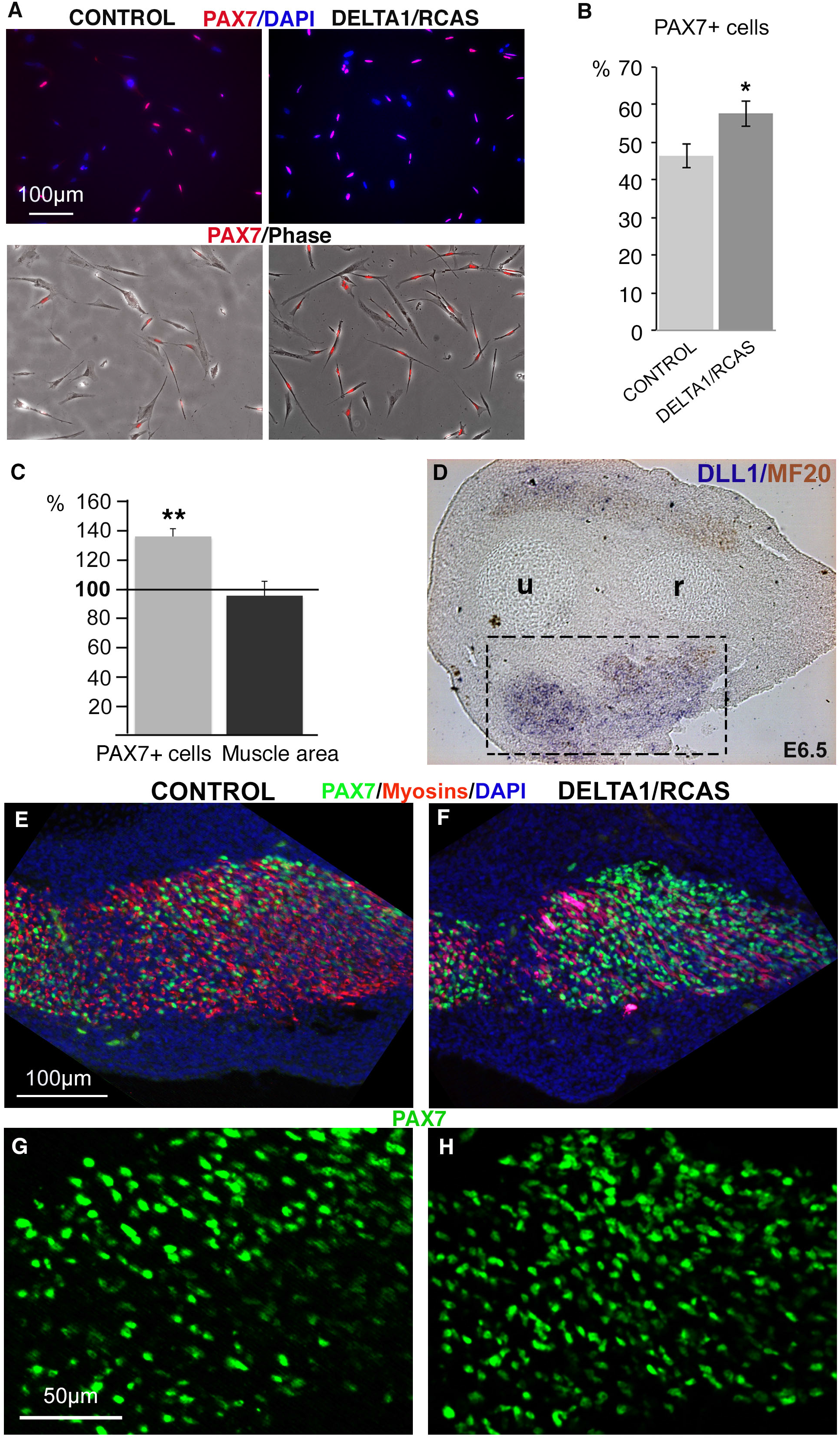
NOTCH gain-of-function experiments increase the number of PAX7+ cells in myoblasts and chicken limbs. (A) Representative fields of chicken foetal myoblasts infected with Empty/RCAS (control) or with DELTA1/RCAS cultured in proliferation conditions, showing merged pictures of PAX7/DAPI or PAX7/phase contrast. (B) DELTA1/RCAS-infected myoblasts displayed a significant increase in the number of PAX7+ cells compared to controls (n=7). (C-H) DELTA1/RCAS-expressing cells were grafted into the presumptive right forelimb buds of E4.5 chicken embryos. (C) Quantification of the PAX7+ cell number in muscle regions of DELTA1-grafted right and control left forelimbs from E6.5 chicken embryos (n=3). DELTA1-grafted right (D,F,H) and control left (E,G) forelimbs from E6.5 chicken embryos were cut transversely and analysed for *DLL1* transcripts (D) and immunostained with the PAX7 and MF20 (myosins) antibodies (E-H). Error bars indicate standard error of the mean for (B) and standard deviation for (C). * p<0.05; ** p<0.01;.

**Supplementary Table 1.**
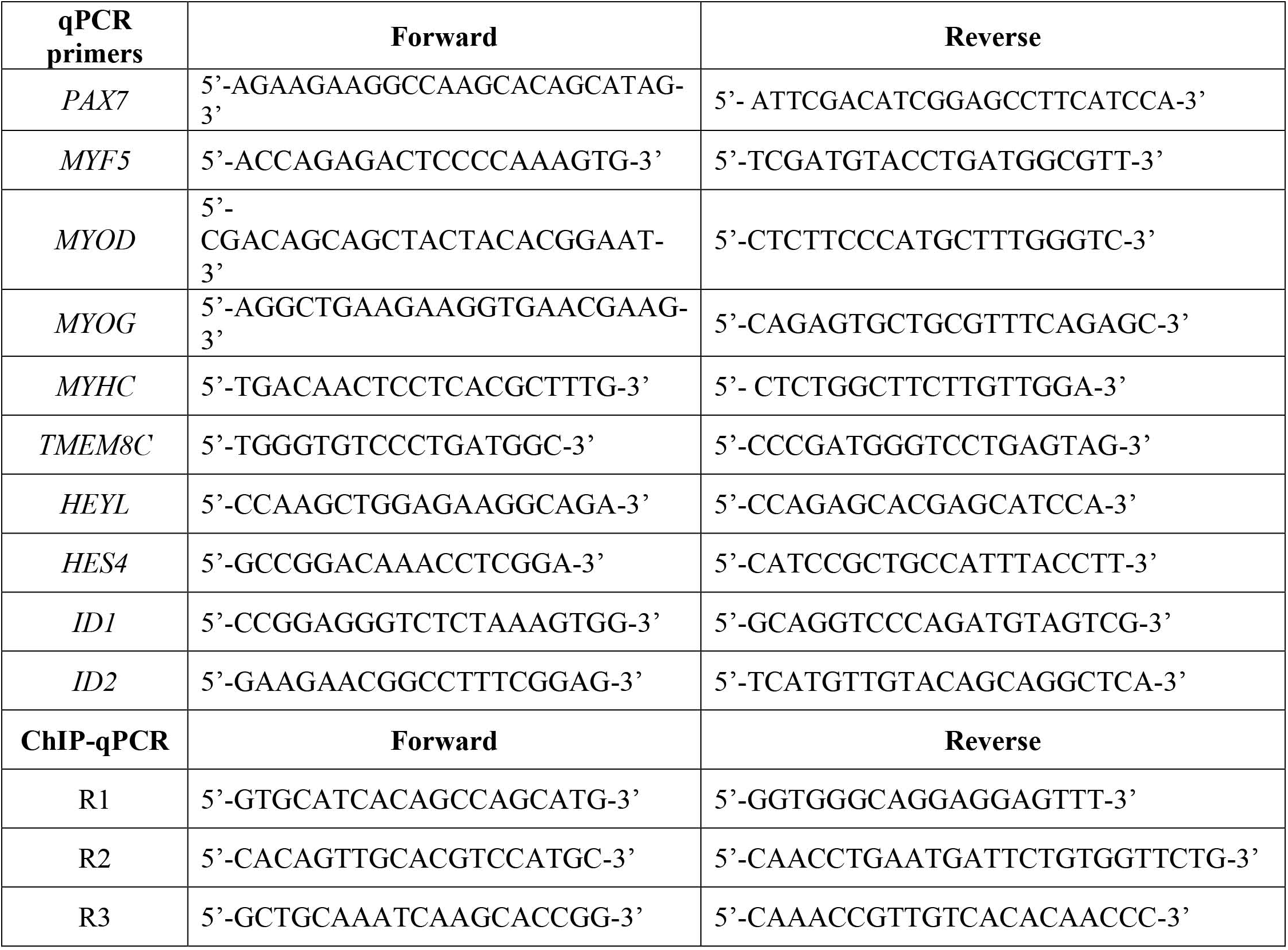
List of primers used in this study.

